# Maturation of cortical endoplasmic reticulum clusters in the mouse oocyte: changes at fertilization

**DOI:** 10.1101/2022.02.28.482320

**Authors:** Huizhen Wang, Lane K. Christenson, William H. Kinsey

## Abstract

Oocytes from many invertebrate and vertebrate species exhibit unique endoplasmic reticulum specializations (cortical ER clusters) thought be essential for egg activation. In examination of cortical ER clusters, we observed they were tethered to previously unreported fenestrae within the cortical actin layer. Further, studies demonstrated sperm preferentially bind to plasma membrane overlying the fenestrae, establishing close proximity to underlying ER clusters. Moreover, following sperm-oocyte fusion, cortical ER clusters undergo a previously unrecognized global maturational change in volume, shape, and calreticulin content that persists through sperm incorporation, before dispersing at the pronuclear stage. These changes did not occur in oocytes from females mated with *Izumo1* -/- males demonstrating that gamete fusion plays an important role in ER cluster maturation. In addition to these global changes seen at sites distant to the sperm, highly localized ER modifications were noted at the sperm binding site as cortical ER clusters surround the sperm head during incorporation, then form a diffuse cloud surrounding the decondensing sperm nucleus. This study provides the first evidence that cortical ER clusters interact with the fertilizing sperm, indirectly through a previous unknown lattice work of actin fenestrae, then directly during sperm incorporation. These observations raise the possibility that oocyte ER cluster-sperm interactions provide a competitive advantage to the oocyte, which may not occur during assisted reproductive technologies such as intracytoplasmic sperm injection.

**Summary Statement:** Sperm-oocyte interactions stimulate global changes in cortical endoplasmic reticulum cluster structure as well as localized responses at the sperm binding site.

## Introduction

Fertilization involves a series of interactions between oocyte and sperm that culminate in one or more high amplitude Ca^2+^ transients which trigger egg activation [1, 2]. The combined Ca^2+^ amplitude is critical for egg activation [3, 4] as inadequate Ca^2+^ stimulus is a significant factor in Oocyte Activation Deficiency following *in vitro* reproductive technologies [5-8]. Oocytes have evolved specialized regions of smooth endoplasmic reticulum (SER) which are thought to support Ca^2+^ transients post-fertilization [2]. Cortical ER specializations were originally identified in marine invertebrates by live cell imaging [9-11] and were described as cisternae or patches interconnected by tubular arrays within the oocyte cortex. Vertebrate animals such as frogs, mice, and humans contain similar cortical ER specializations composed of tightly compacted SER tubules referred to as ‘cortical ER clusters’ [12-17]. These clusters form during oocyte maturation and require NLRP5 (Nucleotide-Binding Oligomerization Domain, Leucine Rich Repeat, and Pyrin Domain Containing 5) expression [18], as well as actin-mediated events for proper tethering to the cortex [19]. Fertilization induced changes in ER structure occur in virtually every species examined, and these changes correlate spatially and temporally with the onset of Ca^2+^ waves or oscillations [14, 20], eventually disappearing at the pronuclear (PN) stage [21]. We were surprised in our study of signaling proteins in the egg cortex, to find multiple, previously undescribed structural changes and rearrangements following sperm-egg interaction. Here we demonstrate that cortical ER clusters are positioned at previously unreported fenestrae within the cortical actin layer and that sperm most often bind over a fenestra and its underlying cortical ER cluster. We further show that sperm-oocyte fusion is required for initiation of maturational changes in ER cluster morphology during fertilization and localized remodeling of cortical ER clusters occurs at the sperm-binding site where clusters surround the sperm head as it is drawn into the ooplasm. These observations provide new insight into the close interaction between oocyte cortical ER clusters and the fertilizing sperm and raise new questions as to the importance of sperm-ER cluster interaction in assisted reproductive technologies.

## Methods

### Animal Handling

Animals were housed in a temperature and light cycle-controlled room and experiments were conducted in accordance with the ‘Guide for the Care and use of Laboratory Animals’ (Institute of Laboratory Animal Resources (U.S.) Committee on Care and Use of Laboratory Animals 1996; National Research Council (U.S.) 2011). Experimental procedures were approved by the University of Kansas Medical Center IACUC committee.

### Collection, fixation and immunolabelling of oocytes

B6D2F1 mice were obtained by crossing inbred C57BL/6NHsd females with inbred DBA/2NHsd males as described (Envigio, Indianapolis, IN). The resulting B6D2F1 females 5 weeks of age were stimulated with 5 IU of Pregnant Mares Serum Gonadotropin (Sigma-Aldrich, St. Louis, MO), followed 48 hours later by 5 IU of human Chorionic Gonadotropin (hCG) (Sigma-Aldrich). In order to avoid handling-related changes in cortical ER cluster morphology, this study used oocytes recovered from mated females and fixed *in situ*. Mating was initiated by addition of a male (minimum of 12 weeks of age) and the females were sacrificed at 30 minute intervals between 15.5-17 hours post-hCG for collection of oocytes. Oviducts were removed and submerged in fixative containing 2% formaldehyde with 1% saturated picric acid in PBS. The ampullae were dissected free of remaining oviductal tissue and then cut at one end to release the cumulus-enclosed oocytes into the fixative with a minimum of handling. As reported elsewhere [22], we observed that prolonged formalin fixation improved retention of organelle morphology. Therefore, after one hour incubation at 25°C, the dish containing cumulus enclosed oocytes was transferred to a refrigerator for an additional 20 hours of fixation at 4°C. After fixation, the cumulus enclosed oocytes were washed in Fixation and Handling Medium (Specialty Media Inc., Phillipsburgh, N.J.) containing 4mg/ml BSA (FHM-BSA) (Sigma-Aldrich), then incubated with 10mg/ml hyaluronidase (Sigma-Aldrich) for 10 min. Cumulus-free oocytes were washed in PBS containing 4mg/ml BSA and transferred to Terasaki plates (NUNC corp. Rochester, NY) for immunolabeling. Oocytes were incubated in blocking buffer consisting of PBS including 0.1% saponin (Sigma-Aldrich) and 3mg/ml BSA for 30 minutes, then incubated with primary antibodies diluted 1/100 in blocking buffer overnight. Primary antibodies used include rabbit anti-human calreticulin (CRT; Abcam, Cambridge, UK (AB92516)) as a marker for ER [23] and rat anti-mouse CD9 (BD-Pharmingen, Franklin Lakes, NJ, (553758)) as a plasma membrane marker [24]. Specificity of the CRT antibody was tested by addition of a synthetic peptide AbCam AB180826 to the primary antibody at 1:100 excess (Supplemental Data Fig 1). Secondary labelling was done for 1 hour in blocking buffer containing Alexa Fluor 488 goat anti-rabbit IgG (InVitrogen, Waltham, MA (A11008)) or Alexa Fluor 488 goat anti-rat (Invitrogen, (A11006)), or Alexa Fluor 635 goat anti-rabbit (InVitrogen(A31577)) diluted1:100 in blocking buffer. Alexa Fluor 568-phalloidin (Invitrogen) (2 U/ml), and Hoechst 33342 (0.1mg/ml) were included with all secondary antibodies to label actin and DNA, respectively and labelled oocytes were transferred to glycerol mounting medium (Abcam, AB188804) prior to imaging.

### Live Cell Imaging

Experiments that involved live cell imaging were performed on cumulus-enclosed oocytes recovered from mated females. Cumulus-enclosed oocytes were treated with hyaluronidase (0.1mg/ml) for 10 minutes to remove cumulus cells [25]. The oocytes with sperm visible in the perivitelline space were washed in KSOM (EMD Millipore Corp. Billerica, MA) containing 5mg/ml BSA (KSOM-BSA) then incubated in 500ul of KSOM-BSA containing 1µM ER Tracker-green (Life Technologies Corp. Eugene, OR) for 30 minutes at 37°C and 5% CO2 in a humidified incubator. Then 0.5µl of SiR-actin (Cytoskeleton Inc. Denver, CO) (1mM) was added to achieve a final concentration of 1µM and the oocytes were returned to the incubator for an additional 30 minutes. Oocytes were then washed in KSOM-BSA and transferred to a poly A-lysine (Sigma-Aldrich) treated Delta-TPG plate (Bioptics, Butler, PA) containing a 10ul drop of KSOM-BSA containing Hoechst 33342 (1ug/ml) and covered with pre-equilibrated mineral oil. The plate was mounted in a Chamlide (Quorum Technology, Inc. Guelph, ON, Canada) TC-L stage top environmental chamber to maintain temperature at 37°C and a CO_2_ level of 5%. Live cell imaging was performed with a Nikon A1R confocal microscope which was used to take tangential scans through the cortical region to determine the relationship between cortical ER clusters and actin fenestrae. SiR-actin was imaged with a 640nm laser and ER-tracker green was imaged with a 488nm laser.

### *In vitro* fertilization

In vitro fertilization was performed on zona-free oocytes as described [25] using a low concentration of capacitated sperm (2×10^4^ per ml) and an incubation period of 20 minutes post insemination. Oocytes were then fixed as described above.

### Microscopy and image analysis

Confocal microscopy of fixed samples was performed with a Nikon A1R confocal microscope, using sequential scans with constant beam intensity. Images were recorded from Z-planes at 1µm intervals through the entire oocyte.

Image analysis was performed to determine the maturation state of cortical ER clusters in oocytes recovered from mated females. The General Analysis tool in Nikon Elements Advanced Research Imaging software (5.02.01 build 1270) was used to create a macro to identify maturing cortical ER clusters. A threshold was used to establish a binary defining the oocyte cortex as that region within 2µm of the cortical actin layer. Within the oocyte cortex, objects with CRT fluorescence intensity above 1200 (approximately 2.5 times the CRT fluorescence of ER in the central cytoplasm) and with a diameter greater than 2µm were considered to be ‘maturing ER clusters’. After the General Analysis macro was run on Z-planes from each egg, the binary editor was used to remove cumulus cells and Meiotic chromosome-associated CRT accumulations by hand. Once the 2D binaries within a Z-stack were established, the 3D Object Measurement tool was used to connect them into 3D objects for measurement of matured ER cluster number, volume, and CRT content. STimulated Emission Depletion (STED) microscopy was performed with a Leica TCS SP8X STED ONE microscope.

### Statistics

Quantification of the number of matured cortical ER clusters, their volume and relative CRT content was performed using multiple experimental replicates described in each Table or Figure legend. Samples were found not to exhibit a normal distribution, so comparisons between experimental groups were performed using the non-parametric Kruskal-Wallis Analysis of Variance test with the mean or median values reported +/- SEM. P values less than 0.05 were considered significant.

## Results and Discussion

### Cortical ER clusters are localized at fenestrae in the cortical actin layer

While previous studies have documented migration of ER clusters to the cortex and shown that cortical localization is required for normal Ca^2+^ signaling at fertilization [13, 14, 19], little is known about the process that directs arrangement of the ER clusters at the cortical actin layer and plasma membrane. Using confocal immunofluorescence imaging to define the spatial relationship between ER clusters and the actin layer, we observed oocytes from virgin (non-mated), as well as mated females exhibited a uniform distribution of lamellar ER structures in the central cytoplasm, while ER clusters were concentrated in the oocyte cortex (Fig. 1 A, B, (arrows)). Reconstructed 3-dimensional images revealed that most ER clusters were arranged underneath fenestrae in the cortical actin layer (Fig.1A’, B’). As expected, the cortical ER clusters were absent from the actin cap region overlying the meiotic II chromosomes as previously described [21] (Fig. 1A’,B’(arrows)) and it was evident that actin-layer fenestrae were absent from the actin cap as well (Fig. 1A’’, B’’(arrows)). These fenestrae in the actin layer overlying the ER clusters have, to our knowledge, not been recognized or described previously. Initially concerned that fenestrae may represent a fixation artifact, cell-permeant markers for ER (ER tracker-green) and for f-actin (SiR-actin) were imaged in live oocytes. Tangential z-plane images of live oocytes revealed the presence of f-actin fenestrae (Fig. 1C, C’’ (arrowhead)) that appear similar to those in the fixed oocytes (Fig. 1A’, A’’). Live oocytes also exhibited cortical ER clusters positioned underneath the actin fenestrae (Fig, 1C, C’) confirming the actin fenestrae and associated ER clusters were not fixation artifacts (Fig. 1A, A’). To determine whether fenestrae represented holes in the cortical actin layer, oocytes were labelled with a combination of phalloidin, anti-CRT, and the plasma membrane (PM) marker anti-CD9. Images taken at higher magnification revealed that fenestrae (Fig. 2 (arrows)) retain a thin layer of f-actin positioned close to the PM, with ER clusters arranged in close proximity to this thin actin layer.

**Fig. 1.**
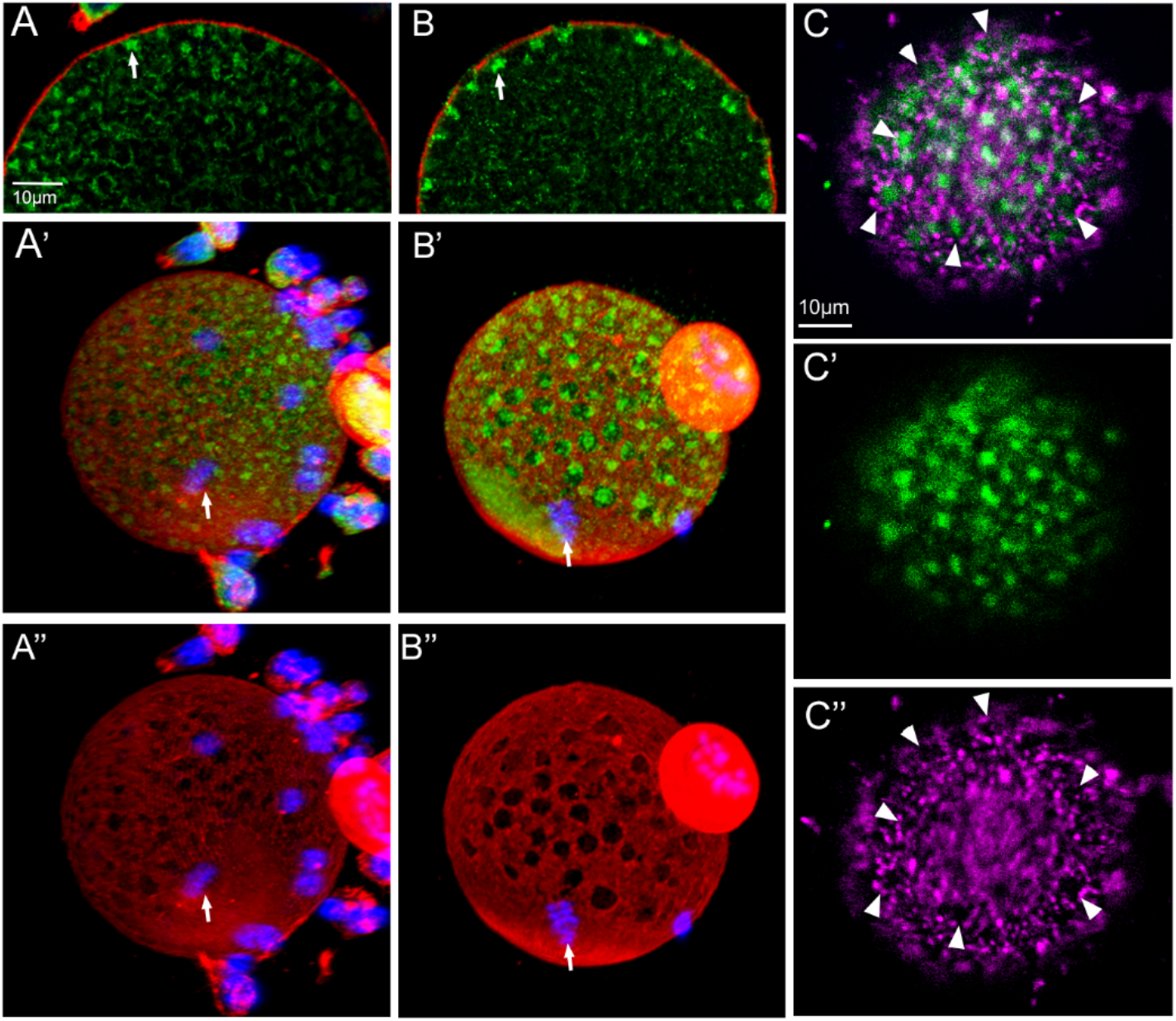
ER clusters are positioned under f-actin fenestrae in fixed and live oocytes. Oocytes from virgin (A) or mated (B) females were fixed and labeled with anti-CRT (green), alexa568-phalloidin (red), and Hoechst 33342 (Blue). Arrows in panels A, B denote cortical ER clusters in an equatorial view. Three-dimensional reconstructions demonstrate f-actin fenestrae (red) and underlying cortical ER clusters (green) with the meiosis II chromosomes (blue) indicated by arrows in panels A’, A’’, B’, B’’. In vivo fertilized oocyte (C, C’ and C’’) imaged live with a confocal microscope, illustrate f-actin fenestrae labelled with SiR-actin (purple) and cortical ER clusters labelled with ER-tracker (green). Panels C’ and C’’ show in the individual green channel and far-red channel, respectively. Arrows in C and C’’ indicate individual fenestrae for reference. Magnification is indicated by the bar which represents 10µm.

**Fig 2.**
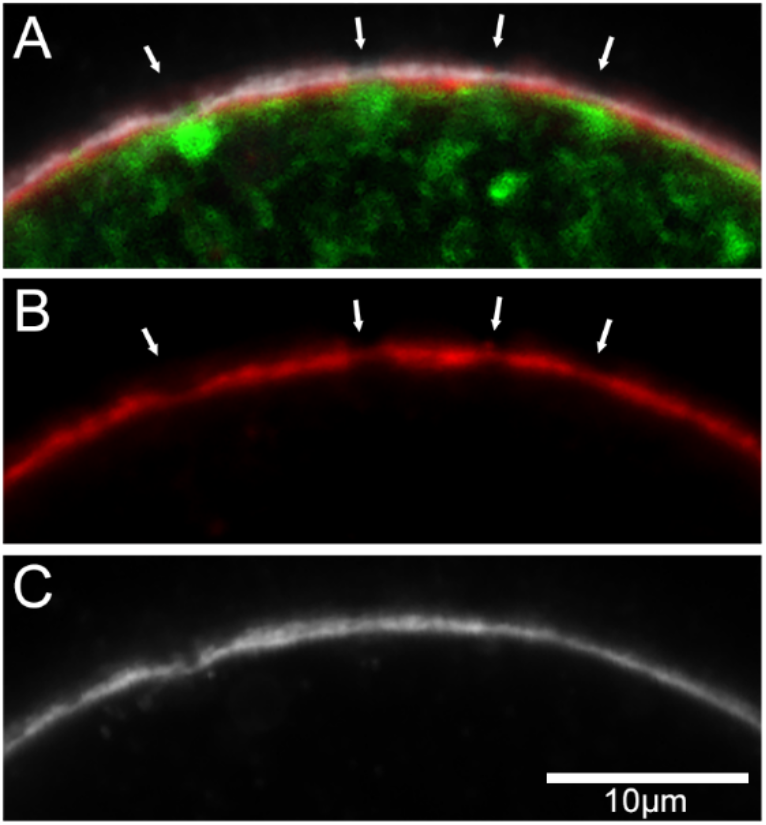
Relationship of oocyte PM, fenestra, and ER clusters. Oocytes were labelled with anti-CRT (green) to detect ER clusters, Alexa Fluor 568-phalloidin (red) to detect f-actin, and anti-CD9 (white) to detect plasma membrane (PM). Composite image of a fertilized oocyte (panel A) showing PM, f-actin, and CRT; panel B shows the f-actin label only with attenuated regions at fenestrae and panel C shows the uninterrupted PM label (anti-CD9) only. Arrows in panel A and B indicate the position of actin layer fenestrae and associated cortical ER clusters relative to the PM. Magnification is indicated by the bar which represents 10µm.

The actin layer fenestrae appear to allow ER clusters to become positioned closer to the PM than would be possible at other (interstitial) regions of the actin layer, which could potentially allow ER clusters to establish contact with the PM. It is also intriguing to speculate that the thinner layer of f-actin inside fenestra may be more easily remodeled than the ‘structural’ regions between fenestrae and could therefore facilitate PM events such as microvillus elongation and ‘egg membrane wave’ formation during fertilization [24]. In any case, actin layer fenestrae likely have unique properties that tether ER clusters, however the question of whether fenestrae attract the ER cluster or the cluster modifies the actin layer to create a fenestra remains unanswered.

### Cortical ER clusters undergo global maturational changes in response to sperm-oocyte interaction

Early live cell analysis of mouse oocytes demonstrated changes in the number and localization of cortical ER clusters in response to fertilization [21]. However, higher resolution analyses obtained through confocal imaging of carefully fixed oocytes [22] revealed a range of structural changes that led us to revisit this process with attention to different progressive stages of sperm-egg interaction [26]. Oocytes from virgin females contained cortical ER clusters approximately 2µm in diameter, that were irregular in shape and contained a diffuse array of CRT-containing ER tubules (Fig. 3A, arrows). In contrast, oocytes from mated females contained morphologically distinct cortical ER clusters with increased volume and CRT content (Fig. 3B, arrows). These ER clusters eventually reached a mature form that were near spherical in shape with more densely packed CRT (Fig. 3C, arrows). Comparison of unfertilized oocytes from virgin females (Fig. 4A) to oocytes with bound or fused sperm (Fig. 4B arrow) demonstrates the initial maturational increase in ER cluster structure. Oocytes fixed while in the process of incorporating sperm (Fig. 4C, C’) demonstrate the substantial increase in CRT content of ER clusters as indicated by increased fluorescence intensity. Oocytes with sperm undergoing nuclear de-condensation exhibited a reduction in the number of mature ER clusters and occasional detachment of the largest ER clusters from the oocyte cortex (Fig. 4D). Finally, during the PN stage the number of matured ER clusters was greatly diminished, leaving instead small granular ER structures which no longer retained association with the cortical actin layer (Fig. 4E; Supplemental data Fig 2B-F).

**Fig. 3.**
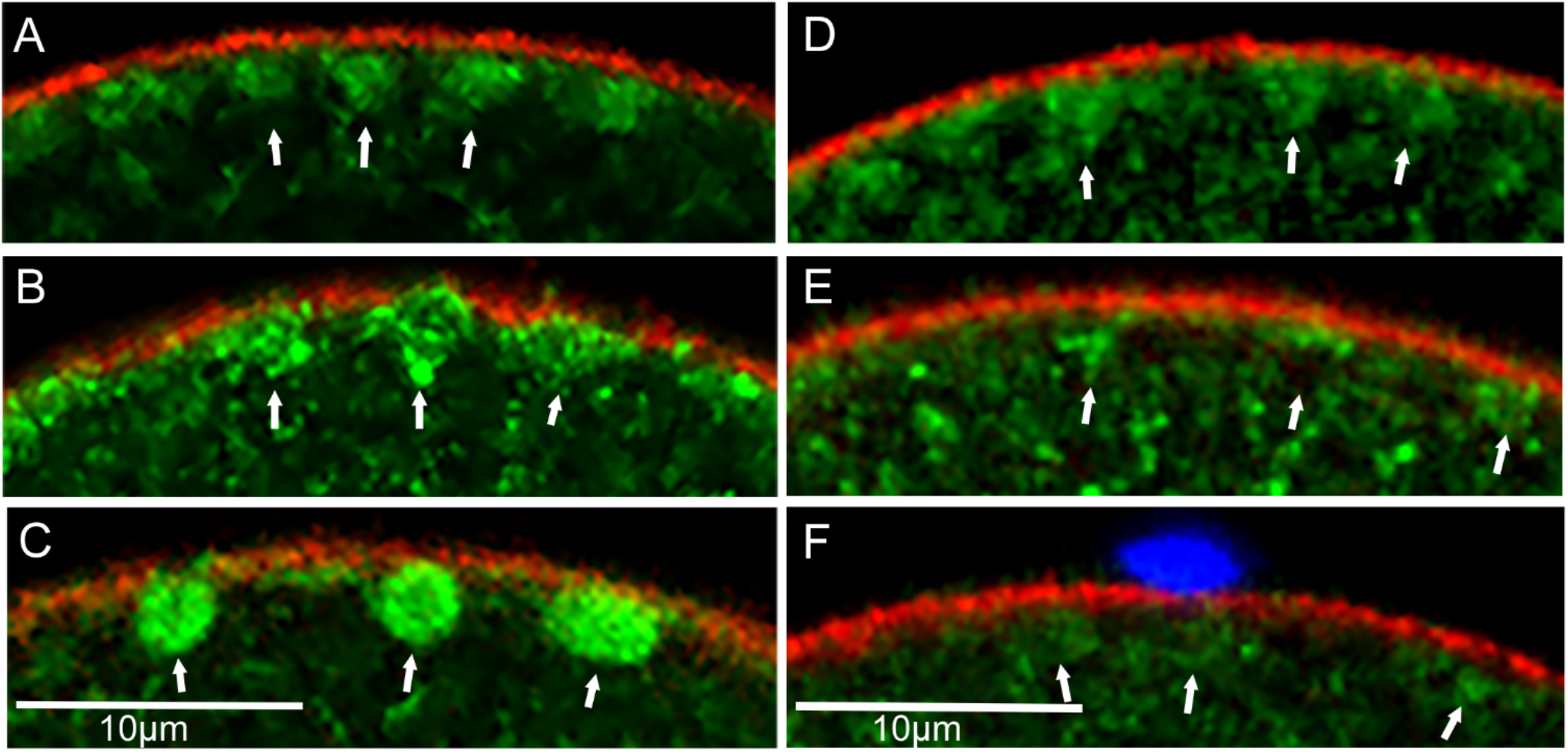
ER cluster morphology changes following mating and requires gamete fusion. Oocytes from virgin (A) females exhibit immature cortical ER clusters, while oocytes from females mated with *wt* males (B, C) exhibit clusters at various stages of maturation. In contrast, oocytes from females mated with Izumo1-/- males (D-F) retain cortical ER clusters in an immature state, similar to virgin (A) females. Arrows indicate the position of clusters. Panel F shows an Izumo1-/- sperm bound to an oocyte. Magnification is indicated by the bars which represent 10µm.

**Fig. 4.**
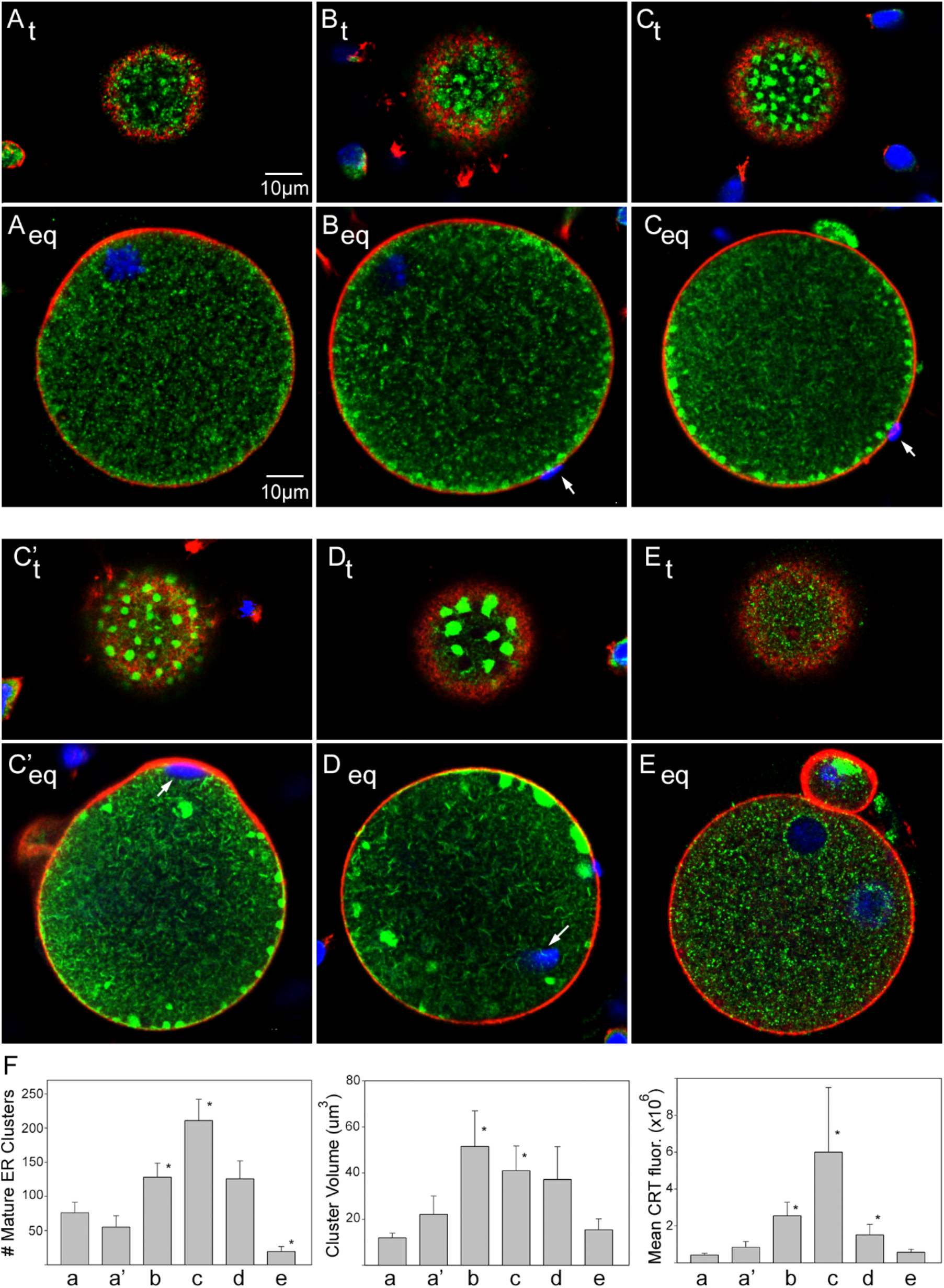
Maturation of cortical ER clusters at progressive stages of sperm-egg interaction. Oocyte images (A-E) include both a tangential (t) and equatorial views (eq) with white arrows denoting sperm, with panel F showing quantitative image analysis results. Stages include: virgin (non-mated) unfertilized oocytes (A), oocyte with bound sperm (B), oocytes with incorporating sperm (C and C’), oocytes with decondensing sperm (D), and oocyte at the PN stage (E). Magnification is indicated by the bar which represents 10µm. Image analysis results (F) shown left to right depict the number of matured cortical ER clusters, mean cluster volume, and mean CRT content in oocytes. Vertical bars (a-e) represent the same stages shown in images above: (a) virgin oocytes (n=22), (b) oocytes with bound/fused sperm (n=17), (c) oocytes with incorporating sperm (n=14), (d) oocytes with decondensing sperm (n=10), (e) early PN stage oocytes (n=17) and (a’) oocytes from mated females but lacking bound sperm (n=11). Data represent the results of five replicate experiments, each using at least two females. All z-planes from each oocyte were examined individually and, where necessary, in 3D format to determine the presence or absence of bound-fused sperm and insure the absence of polyspermy. Values represent the mean +/- SEM and (*) indicates that the mean is significantly different from virgin female oocytes as determined by the Kruskal-Wallis Analysis of Variance test.

To quantify changes in cortical ER cluster maturation at the above stages, oocytes from virgin and mated females were imaged at 1µm z-plane intervals through the entire oocyte and the mean values for ER cluster number, volume, and CRT content across the stages were determined (Fig. 4F). Samples from five experiments were analyzed, each using 2-3 male and female mice. Oocytes collected from virgin (Fig. 4F, a) and from mated females which did not have sperm in the perivitelline space (Fig. 4F, a’) exhibited no significant difference in the number, volume or CRT content of matured ER clusters. However, oocytes with bound/fused sperm (Fig. 4F, b), as well as oocytes with incorporating sperm (Fig. 4F, c) exhibited a significant increase in all three parameters of ER cluster maturation. The number and CRT content of ER clusters began to decline during sperm de-condensation (Fig. 4F, d) and almost completely disappeared as oocytes entered the PN stage (Fig. 4F, e).

Once maturation is initiated, enlargement of cortical ER clusters likely involves translocation or flow of ER components from tubular arrays in the central ooplasm to the cortex as shown with the lipophilic dye DiIC18 [9]. Translocation could involve microtubule-mediated ER reorganization as demonstrated in *Xenopus* oocyte extracts [27] and could contribute to ER cluster maturation in the egg cortex by tethering ER proteins directly to microtubules and associated microtubule motor proteins [28]. Actin remodeling may also play a role [19] in positioning ER Clusters at fenestrae, or in stabilizing ER tubules in a compacted, spherical form which could provide maximal ER in close proximity to the PM overlying fenestrae. Functionally, the increase in ER cluster size and CRT density suggests that maturation could significantly increase the capacity of ER clusters to store Ca^2+^ in the oocyte cortex. This would likely promote Ca^2+^ oscillation amplitude and duration as proposed in the *Xenopus* oocyte [15]. Contact sites between ER and PM have been detected in electron microscopic imaging of *Xenopus* oocytes and were hypothesized to play a role in Ca^2+^ uptake during egg activation [29]. Our observation that ER clusters express tubules that extend toward the PM (below) suggests the possibility of ER-PM contact in the mouse oocyte which would support the above hypothesis. Further imaging of mouse oocytes by super-resolution microscopy or by electron microscopy might identify tethering proteins typical of “functional membrane contact sites” [30] and provide insight into how ER clusters function during fertilization.

### Role of gamete fusion

The fact that global maturation of cortical ER clusters was first observed when sperm interacted with the oocyte plasma membrane raised the question of whether sperm-oocyte binding, or gamete fusion provided the stimulus that initiates ER cluster maturation. To test the hypothesis that gamete fusion is required for ER cluster maturation, we took advantage of the fact that *Izumo1*-null sperm can bind to the oocyte PM, but are unable to fuse with oocytes [31]. Oocytes recovered from females mated to *wt* or *Izumo1*^*-/-*^ males were fixed and labelled for detection of CRT and f-actin to determine the number of matured cortical ER clusters per oocyte (Table 1, Fig. 3D-F, arrows denote ER clusters). Oocytes that had fused with sperm were identified by the presence of at least one sperm that had partially or completely penetrated the cortical actin layer. The frequency of sperm-egg fusion was much higher in oocytes from females mated with *wt* males (94% fusion) than in oocytes from females mated with *Izumo1*^*-/-*^ males (0% fusion). The number of mature cortical ER clusters was also dramatically higher in oocytes from females mated with *wt* males compared to those mated with *Izumo1*^*-/-*^ males, demonstrating that ER cluster maturation was highly associated with gamete fusion. While the signal(s) that initiate the maturational response of ER clusters remain unknown, our observation that oocytes from females mated with *Izumo1*^-/-^ males failed to undergo ER cluster maturation demonstrates that sperm-egg fusion is a prerequisite to ER cluster maturation. Upstream events that may induce ER cluster maturation include release of PLCζ at the sperm fusion site followed by calcium-induced calcium release that could propagate along interconnecting cortical ER clusters triggering global cluster maturation. Other possibilities include reorganization of the cortical actin layer in response to the initial Ca^2+^ transient [32] which involves Src-family [33] or other protein kinases such as Focal Adhesion Kinase (PTK2A) [34], which, in other systems, can co-localize with ER and can complex with IP3r1 to spatially localize Ca^2+^ signaling [35].

**Table 1.** Cortical ER cluster maturation requires fusion-competent sperm.

| 3 genotype | Oocytes (n) | % Fusion | Mature ER Clusters/<br>4 Oocyte |
| --- | --- | --- | --- |
| 5 *wt* | 17 | 94 | 137 +/- 16 |
| 6 *Izumo1*<sup>-/-</sup> | 27 | 0 | 19 +/- 9 * |

7 Oocytes recovered from females mated to *wt* or *Izumo1*<sup>-/-</sup> males were fixed, labeled with anti-CRT, and imaged as 8 described in Materials and Methods. To minimize possible genotype-dependent differences in zona-penetration, only 9 oocytes with sperm in the perivitelline space or in the ooplasm were analyzed. Oocytes with a sperm that had partially or completely penetrated the cortical actin layer during the 15.5-17 hr post-hCG collection period were judged to have undergone gamete fusion. The number of mature ER clusters/oocyte in the *wt* and *Izumo1*<sup>-/-</sup> groups was compared using the Kruskal-Wallis One Way Analysis of Variance on Ranks test and the median number of mature clusters/oocyte is indicated +/- SEM. (\*) indicates there is a significant difference in median values (P<0.001). Experiments were replicated three times using 2 breeding pairs during each replicate.

### Sperm binding at fenestrae and interaction with cortical ER clusters

Observation of oocytes in multiple experiments suggested that sperm frequently bound to regions of oocyte PM overlying a cortical ER cluster (Fig. 5, A-F). Quantification revealed that 81% (Table 2) of bound sperm were positioned over a cortical ER cluster even though clusters only occupy 20-30% of the oocyte surface, which led us to look for possible interactions between the sperm and cortical ER clusters. The observation that most sperm binding sites are positioned at regions of oocyte PM overlying a cortical ER cluster raises the possibility that the PM overlying an actin fenestra/ER cluster may be specialized to act as a high affinity sperm binding site [24, 36]. While consistent with the observations, proof would require demonstration that the PM overlying cortical ER clusters is enriched in sperm binding proteins and ER tethering proteins. We also observed that cortical ER clusters positioned underneath a bound sperm often exhibited morphological changes not seen in ER clusters distant from the sperm. A common change in ER cluster shape included the extension of multiple CRT-containing tubules from the ER cluster toward the sperm head (Fig. 5A, B). These may represent ER tubules within enlarged microvilli typically seen at the sperm binding site [37], or they may represent an ER tubule traversing a fusion pore to establish close proximity to a fused sperm. Another change observed at sperm binding sites involved transformation of an ER cluster into an annulus underneath the bound sperm (Fig. 5B, E), some of which were associated with a central cavity devoid of CRT (Fig. 5 C). Regardless of mechanism, binding of a sperm at a cortical ER cluster seems to trigger an active response by the ER since, using STED microscopy, we observed CRT-rich ER tubules extending from the ER cluster toward the bound/fused sperm (Fig. 5D, E). During sperm incorporation, sperm could be seen penetrating through the cortical actin layer where the sperm head established close proximity to a cortical ER cluster (Fig. 5F). As sperm incorporation proceeded, cortical ER clusters often formed a collar-like annulus surrounding the sperm head (Fig. 5G,G’,H,H’ yellow arrows). In addition, smaller ER structures were observed to surround the incorporated sperm head (Fig. 5H’ white arrows). Upon full incorporation of the sperm into the ooplasm, the sperm chromatin began to de-condense and ER clusters lost their compact structure (Fig. 5I). ER tubules then formed a diffuse cloud surrounding the sperm once it was no longer associated with the cortical actin layer (Fig. 5J).

**Fig. 5.**
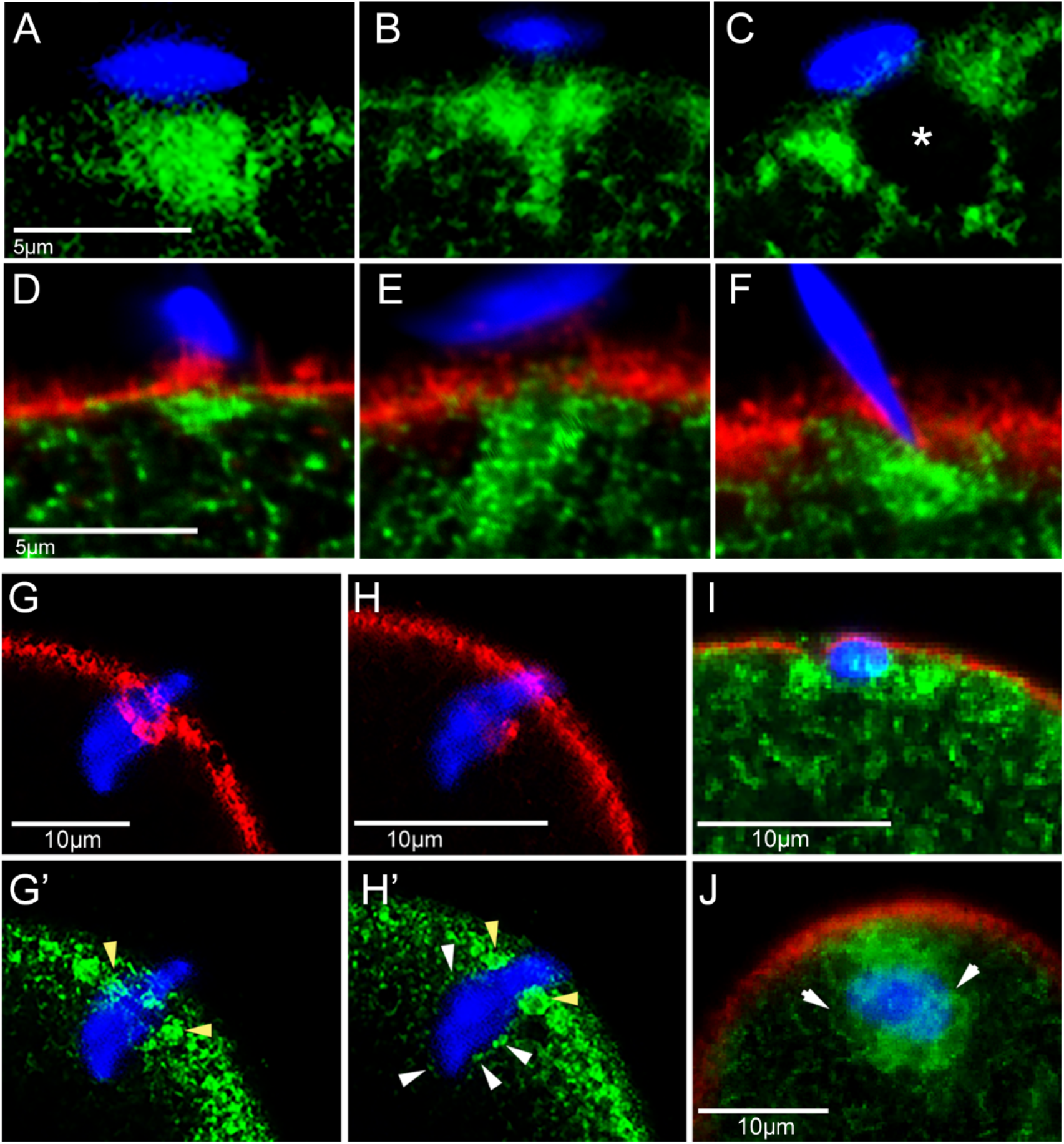
Cortical ER clusters at the sperm binding site and during and after sperm incorporation. Confocal (A-C, G-J) and super-resolution (D-F) microscopic images of sperm (blue), oocyte cortical ER (green), and f-actin (red). Panel A shows a maturing ER cluster with CRT-rich tubules in close proximity to the sperm PM, Panel B shows a maturing funnel-shaped ER cluster with CRT-rich tubules in close proximity to the sperm PM, while Panel C shows a CRT-cluster with enclosed cavity (*). Panels D-F include images obtained by super-resolution microscopy to demonstrate f-actin-containing cell processes extending from the oocyte to establish close proximity to the sperm PM. Panel F shows an example of a sperm head penetrating the oocyte actin layer to establish close proximity with an ER cluster. Panels G, G’-H, H’ show ER clusters and f-actin forming a collar or annulus around a sperm during incorporation (yellow arrowheads), and other ER structures (white arrowheads) enveloping the sperm head. Panel I shows a later stage where the sperm head has been incorporated and the annulus has begun to dissipate. Panel J represents a later stage showing an oocyte ER ‘cloud’ surrounding an incorporated sperm head. Magnification is indicated by the bar which represents 5 or 10µm.

**Table 2.** Sperm binding at cortical ER cluster sites.

|  |  |  |  |  |  |
| --- | --- | --- | --- | --- | --- |
| 0 | Source | # oocytes with | # binding | # bound at | % bound |
| 1 |  | bound sperm | sites | ER Cluster | at ER clusters |
| 2 | Mated | 26 | 28 | 23 | 81.4 +/- 9.2 |
| 3 | IVF | 30 | 252 | 198 | 80.0 +/- 5.1 |

4 Oocytes recovered from mated females or from virgin females processed through in vitro fertilization (IVF) were fixed, 5 labeled with anti-CRT, and imaged as described in Materials and Methods. Sperm bound to the oocyte PM were 6 classified as either 'bound' (at least partially over an ER cluster), or 'not bound' (not over an ER cluster). Experiments 7 using mated females were replicated eight times using a total of 11 females and IVF experiments were replicated seven 8 times using 7 females. The mean % of sperm that were bound over an ER cluster is indicated +/- SEM.

There are reasons to suspect a theoretical advantage in having an ER cluster close to the fertilizing sperm. One potential benefit of this arrangement would be that the ER cluster serves as a Ca^2+^ reservoir to facilitate membrane fusion, or to promote early actin remodeling events localized to the sperm binding site during the initial steps of sperm incorporation. Additionally, an ER cluster physically associated with the fertilizing sperm might be able to respond more quickly to PLCζ release and thus initiate the first Ca^2+^ transient earlier than would cortical ER clusters located farther away.

In summary, this study presents several novel discoveries involving cortical ER clusters in oocytes. The finding that cortical ER clusters become localized at specialized fenestrae within the cortical actin layer, and that sperm preferentially bind to PM overlying fenestrae suggest that fenestrae play a role in ER cluster function and possibly sperm-egg interaction. The finding that ER clusters undergo structural changes in response to sperm-egg fusion suggests active remodeling of the ER which must involve a control mechanism that deserves further study. In addition to these global events, localized ER cluster remodeling at sperm binding site suggests the presence of unknown interactions between the sperm and the ER cluster which may promote the early stage of Ca^2+^ oscillations and involve other unknown control mechanisms.

## Acknowledgements

We acknowledge assistance from Sarah Tague for help with image analysis.

## Competing Financial Interests

The authors declare no competing interests relating to the manuscript.

## Funding

This work was supported by NICHD-HD062860, as well as a University of Kansas School of Medicine Investigator Assistance Award, and utilized core facilities supported by NICHD-HD02528 and HD090216.

## Data Availability

Images used for the figures in this study are available at “http://www.cellimagelibrary.org “. Datasets used in this study are available from the corresponding author on reasonable request.

## Supplemental Data

**Supplemental Data Fig. 1.**
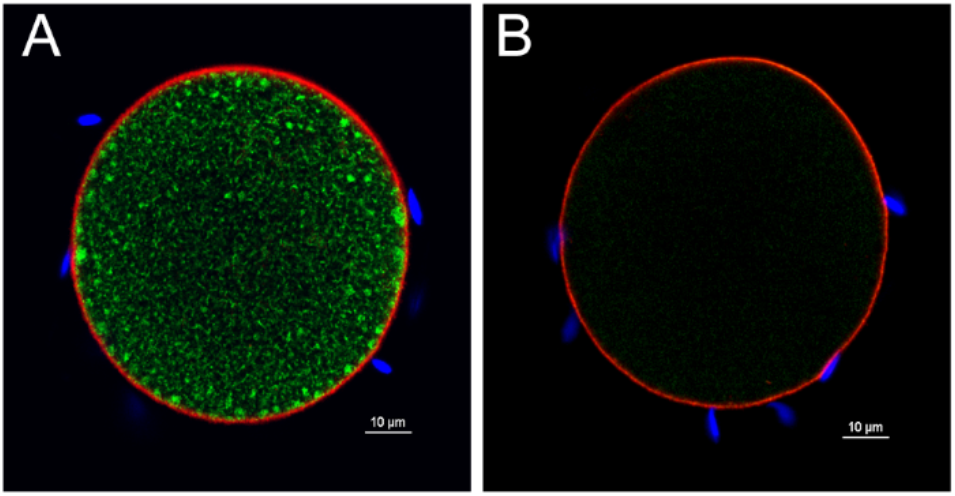
Specificity of anti-CRT antibody. (A) Fertilized oocytes were fixed and labelled with the primary anti-CRT antibody, alexa 568-phalloidin (red), and Hoechst 33342 (blue) as described in ‘Methods’. (B) Another group of oocytes was labelled identically except that a synthetic peptide (Abcam AB180826) duplicating the CRT antigen to which the primary antibody was produced, was included as a competitive inhibitor at 100ug/ml. After washing, the oocytes were incubated with secondary antibody to label CRT (green). The peptide almost completely blocked anti-CRT binding demonstrating the specificity of the antibody. Magnification is indicated by the bar which represents 10µm.

**Supplemental Data Fig. 2.**
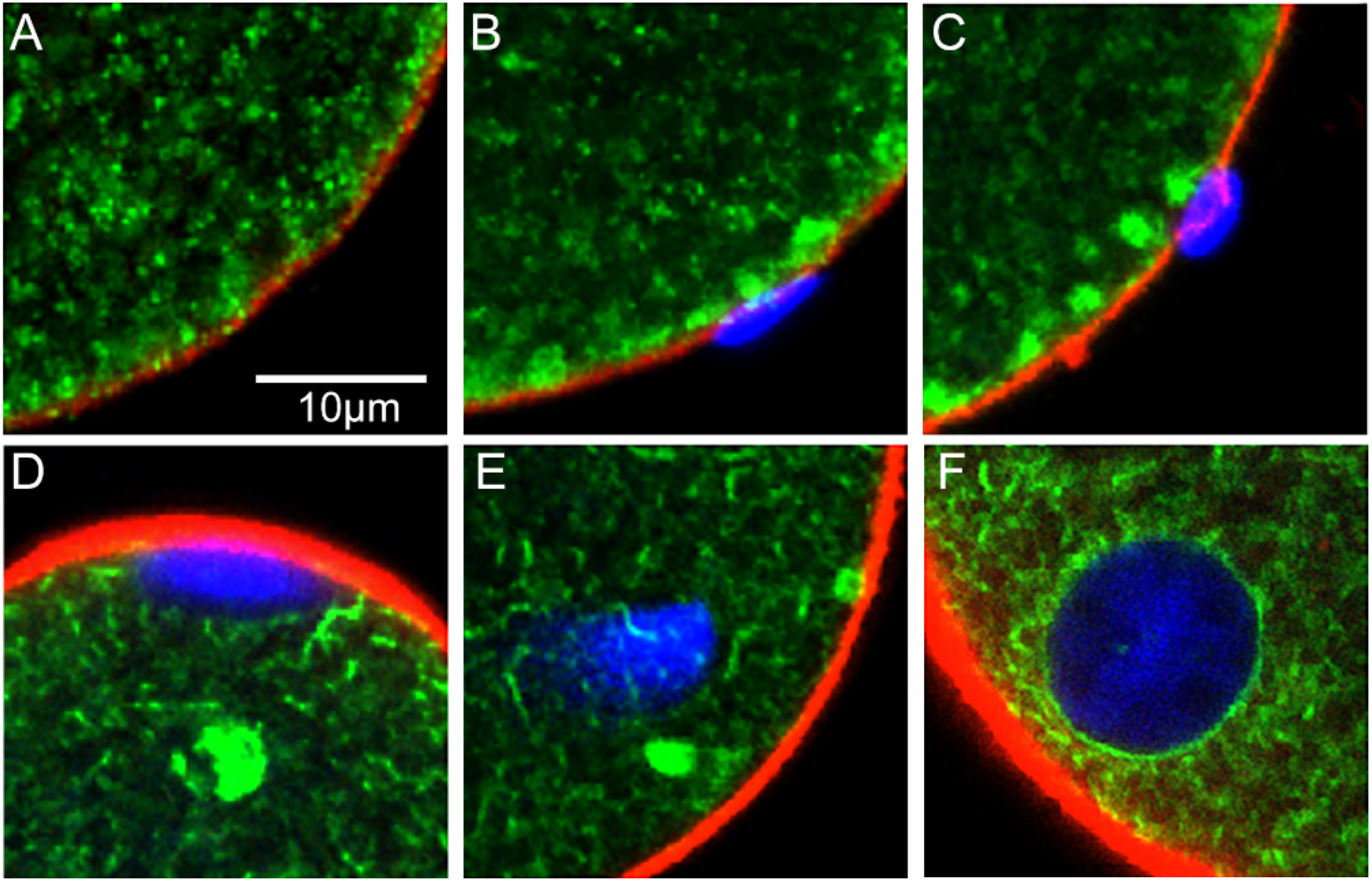
Enlargement of regions of interest from Fig. 4. These images are enlargements of regions of oocytes in Fig. 4. (A) Typical region of oocyte cortex in an unfertilized oocyte with immature clusters, (B) bound sperm and adjacent maturing ER clusters, (C) early sperm incorporation with maturing ER clusters, (D) early sperm de-condensation, (E) late sperm de-condensation, (F) early PN stage showing CRT concentrated at nuclear envelope. Magnification is indicated by the bar which represents 10µm.

## Notes

### Competing Interest Statement

The authors have declared no competing interest.

